# Spatially-divergent metabolic impact of experimental toxoplasmosis: immunological and microbial correlates

**DOI:** 10.1101/2025.01.13.632649

**Authors:** Mahbobeh Lesani, Caitlyn E. Middleton, Tzu-Yu Feng, Jan Carlos Urban Arroyo, Eli Casarez, Sarah E. Ewald, Laura-Isobel McCall

## Abstract

Maladaptive host metabolic responses to infection are emerging as major determinants of infectious disease pathogenesis. However, the factors regulating these metabolic changes within tissues remain poorly understood. In this study, we used toxoplasmosis, as a prototypical example of a disease regulated by strong type I immune responses, to assess the relative roles of local parasite burden, local tissue inflammation and the microbiome in shaping local tissue metabolism during acute and chronic infection. Toxoplasmosis is a zoonotic disease caused by the parasite *Toxoplasma gondii*. This protozoan infects the small intestine and then disseminates to nearly every organ in the acute stage of infection, before establishing chronic infection in the skeletal muscle, cardiac muscle and brain. We compared metabolism in eleven sampling sites in C57BL/6 mice during the acute and chronic stage of *T. gondii* infection. Strikingly, significant metabolic changes were observed in the large intestine and colon during chronic infection, organs not associated with *T. gondii* persistence. Overall, major spatial mismatches were observed between metabolic perturbation and local parasite burden for both disease stages. In contrast, a stronger association with indicators of type I immune responses was observed, indicating a tighter relationship between metabolic perturbation and local immunity, than with local parasite burden. In addition, we observed significant changes in microbiota composition with infection, and candidate microbial origins for multiple metabolites impacted by infection. These findings highlight the metabolic consequences of toxoplasmosis across different organs, and their regulators.

**Importance:** Inflammation is a major driver of tissue perturbation. However, the signals driving these changes on a tissue-intrinsic and molecular level are poorly understood. This study evaluated tissue-specific metabolic perturbations across eleven sampling sites following systemic murine infection with the parasite *Toxoplasma gondii*. Results revealed relationships between differential metabolite enrichment and variables including inflammatory signals, pathogen burden and commensal microbial communities. These data will inform hypotheses about the signals driving specific metabolic adaptation in acute and chronic protozoan infection, with broader implications for infection and inflammation in general.

## Introduction

Toxoplasmosis is a zoonotic disease caused by *T. gondii* parasites, an organism that is estimated to infect more than 30% of the world population. Of the two billion people infected by *T. gondii*, Latin America, Africa, Eastern/ Central Europe, and Southeast Asia have the highest infection rate (1). Once infected, patients often harbor *T. gondii* for life. *T. gondii* infection is controlled by a canonical Type I CD8+ T cell response (2). Immunocompetent patients have mild, flu-like symptoms that are often undiagnosed at the time of infection. Immune-compromised patients, including people with HIV with low number of T cells and patients receiving immunosuppressive therapies for cancer or organ transplants, are susceptible to primary infection and to the recrudescence of chronic infection leading to *Toxoplasma* encephalitis and uncontrolled systemic infection if treatment is not provided (3). One recent study estimated that more than 13 million people living with HIV are co-infected with *T. gondii* worldwide (4). In immunocompetent mothers, primary infection during pregnancy can lead to congenital toxoplasmosis causing miscarriage, stillbirth, or chorioretinitis, hydrocephalus, and intracranial calcifications (3). An estimated 190,000 new cases of congenital toxoplasmosis occur annually, in addition to the 1.2 million existing ones (5).

*T. gondii* has the remarkable ability to infect most nucleated cell types. The parasite initially invades the small intestine, infecting stromal cells and infiltrating immune cells, which play dual roles, both mediating systemic infection and clearing parasites. IFN-γ plays a critical role in initiating immune clearance of *T. gondii* and polarizing the protective adaptive immune response, which peaks around 14 days after infection. The parasite has developed strategies to evade sterilizing immunity, converting from the rapidly dividing tachyzoite phenotype that dominates acute infection to slowly dividing bradyzoites which form long-lived tissue cysts in the brain, cardiac and skeletal muscle as well as in visceral organs, including the lungs, liver, and kidneys at lower frequency (6).

After infection, the homeostatic balance of glycolysis, glutaminolysis, and fatty acid oxidation shifts to activate immune cells and provide substrates for defense mechanisms against pathogens (7)(8). There is a growing appreciation that metabolism is altered in acute and chronic *Toxoplasma* infection. Studies in BALB/c mice infected with *T. gondii* showed higher metabolic perturbation in the acute stage compared to the chronic stage in the liver (9), brain (10) and spleen (11). Liver and brain cholesterol levels were reduced during acute *T. gondii* infection in Swiss Webster mice, but normalized during chronic infection (*12*). Chronic infection with *T. gondii* can result in the progressive muscle wasting disease called cachexia (13, 14)(15). This has been associated with increases in some serum sphingolipid and glycerophospholipid species and decreased levels of a number of TCA cycle intermediates (16). Previous studies have shown that metabolic perturbation can differ between organs such as the brain (17), serum (10), spleen (11), and liver (9). These studies suggest that metabolic perturbations may depend on location (eg. organ/tissue type) and time point (acute versus chronic stages of infection), parameters linked to shifts in parasite abundance. However, a study directly comparing these parameters has not previously been performed.

To understand the local drivers of host metabolic alterations during *T. gondii* infection, we assessed eleven organs during the acute and chronic stages of *T. gondii* infection. Unexpectedly, the cecum, which does not directly interface with *T. gondii,* had a significantly altered metabolic profile in acute infection, which was sustained during chronic infection. This corresponded with sustained shifts in commensal microorganisms. We found the tightest association between metabolic perturbations and immune responses, whereas no clear spatial association with local tissue parasite burden was observed. Most of the perturbed metabolites were organ-specific, and metabolic changes in each organ were usually infectious stage-specific. Despite this, carboxylic acids, fatty acyls, glycerophospholipids, and organonitrogen compounds were commonly perturbed between organs and stages. These results help us understand the microbial and inflammatory drivers of organ-specific metabolic responses to parasitic diseases over time. In the long-term, this work will enable the identification of metabolic pathways that can be modulated to improve disease symptoms without compromising parasite clearance, consistent with a disease tolerance framework (18).

## Materials and Methods

### Mice

C57BL/6 male mice were purchased from the Jackson Laboratory. Mice were housed in the University of Virginia animal facility for two weeks prior to infection and bedding was mixed to help homogenize the microbiome and minimize cage effects. Mice were infected and monitored in accordance with the University of Virginia Institutional Animal Care and Use Committee, AAALAC, and IACUC protocol #4107.

### Infections and food intake monitoring

Per oral infections were conducted with 25 *T. gondii* cysts of the Type II Me49 strain stably expressing green fluorescent protein (GFP) and luciferase as previously described (19). Briefly, cysts were harvested from chronically infected CBA/J mice, mashed through a 50-micron sieve, and stained with *Dolicos biflorus* agglutinin conjugated to rhodamine (Vector Laboratories). Cysts were counted at 20x magnification based on GFP, rhodamine signal and morphology. Experimental mice were fasted overnight and infected by pipetting into the mouth. Mice were housed on wood chip bedding. Mice were weighed daily and monitored for moribund behavior. Experiments were set up with groups of 5 mice per infection and repeated twice (10 total per condition). The experimental N value for acute infection is (N=9) because 1 mouse did not sera convert (not infected) and for chronic infection (N=7) due to animals meeting euthenasia criteria before the experimental endpoint.

### Tissue harvest for metabolomics

Mice were perfused by intracardiac injection with 10 mL of PBS. 30–50 mg of each tissue was harvested as follows. For the small intestine, cecum and large intestine, contents were gently eliminated from the tissue with curved forceps and a sample of the slurry was isolated. The cecum, large intestine, duodenum and the central portion of the jejunum and ileum segments were isolated. A segment of the right lobe of the liver, the apex and ventricular region of the heart and a quadricep were collected. Samples were flash-frozen in liquid nitrogen using pre-weighed tubes and tissue mass was determined.

### Metabolite extraction

Metabolite extraction was performed as described in Want *et al* (*20*). We have previously validated this method in a variety of infectious disease models (21–24), so that its implementation here enables cross-study comparison. Tissues were resuspended at 50 mg per 175 μL of chilled LC-MS–grade water (Fisher Optima), on ice. Tissues were homogenized by bead beating with a 5-mm steel ball in a pre-chilled TissueLyser (Qiagen) at 25 Hz for 3 min. A fraction of homogenate was set aside and frozen for DNA extraction. For aqueous metabolite extraction, LC-MS–grade methanol (Fisher Optima) spiked with 4 μM sulfachloropyridazine (Sigma-Aldrich) was added to 30 mg (by volume) of homogenized sample to a final concentration of 50% methanol. The sample was homogenized in a chilled TissueLyser (Qiagen) at 25 Hz for 3 min. The homogenate was centrifuged for 15 min at 14,000*g* at 4°C to eliminate debris. The supernatant was dried in a Savant SPD1030 SpeedVac (Thermo Fisher Scientific), sealed and stored at -80. For organic metabolite extraction, the centrifugation pellet was resuspended in 3:1 (by volume) dichloromethane (Fisher Optima): methanol solvent mixture and further homogenized at 25 Hz for 5 min. Samples were centrifuged at 14,000*g* for 2 min at 4°C to eliminate debris. The second centrifuged supernatant was collected and air-dried in the fume hood, sealed and stored at −80°C.

### Liquid chromatography-tandem mass spectrometry data acquisition

All LC-MS analysis methods were previously optimized for tissue small molecule characterization (23, 25). Organic and aqueous extractions were merged by resuspending in 150 μL of 50% MeOH spiked with 2 μM sulfadimethoxine and transferred to a new plate after sonication and centrifugation (5 min, 14,800 RPM). The samples were separated through a C8 LC column (Phenomenex; 0.7 μm, 50 mm × 2.1 mm, 100 Å Kinetex) with a C8 guard cartridge, followed by LC-MS/MS acquisition in positive mode (Thermo Scientific Vanquish UHPLC system and Q Exactive Plus MS). Instrumental performance was verified by acquiring data from a standard mix of 6 known small molecules. The 2994 highest abundance *m/z* found in blank samples and the 6 *m/z* from our standard mix of known molecules were added to the method as an exclusion list. Instrumental methods were as in (26).

### Liquid chromatography-tandem mass spectrometry data analysis

Collected raw data were converted to mzXML with MSConvert software (version 3.0.19014-f9d5b8a3b, Proteowizard, Palo Alto, Santa Clara, CA, USA) (27). mzXML data were processed through MZmine 2 (version 2.53) to generate the feature table (28). All MZmine data analysis parameters are in **Table S1**. Due to observing a batch effect (from changing the column during the run due to high backpressure and changes in peak shape), WaveICA batch removal method was applied to normalize peak area (parameters: alpha=1, cut-off=0.1, K=10). In recent comparative work, WaveICA outperformed all but one batch normalization method on a large dataset (29). PCoA plots (Bray–Curtis dissimilarity metric), PERMANOVA and distance analysis to detect differences between groups were done through QIIME2 (30)(31). To annotate the features and visualize annotation mirror plots, GNPS (Global Natural Products Social molecular networking) was used (32)(33), and annotations retrieved using an automated code (22). All annotations were visually inspected for match quality, and for biological plausibility, as described in Theodoris et al (34). All annotations are at confidence level 2/3, according to the metabolomics standards initiative (35). GNPS parameters were as in (36). MolNetInhancer was used to classify features into chemical families (37) using ClassyFire chemical ontology (38). To visualize the features and their structural relationship as determined by molecular networking, Cystoscape (version 3.9.0) was used (39). Random forest classification analysis (40) was used to find the metabolites that were perturbed the most during infection at different stages (1000 trees). UpSet plots were generated through https://asntech.shinyapps.io/intervene/. R and python codes (https://upsetplot.readthedocs.io/en/stable/) were implemented to show common and specific perturbed metabolites related to each organ. 3D ‘ili models (https://ili.embl.de/) were used to generate 3D models of metabolome perturbation and parasite burden in different organs (41). The 3D model was adapted from the model built in Hossain et al (25) to add heart, intestinal contents, liver and quadriceps, using Meshmixer, as described in Dean et al (24). Undetermined values (below limit of detection by qPCR) for parasite burden were substituted with the lowest number of parasites detected in each organ divided by five. Median distance correlation plots were generated using ggplot2 (version 3.5.1), with linear regression R^2^ and equations determined in Excel.

### Assessment of parasite burden

DNA was isolated from 20 mg of tissue homogenate in water, generated as described above. Genomic DNA was isolated with DNeasy Blood & Tissue Kit (Qiagen# 69506/69581) per manufacturer’s instructions. Parasite burden was measured by quantitative RT-PCR using 100 ng DNA per reaction on a QuantStudio 6Flex (Applied Biosystems) using Thermo Fisher (cat # 4444557) TaqMan Fast Advanced Master Mix as previously described (13). The following Taqman primer/probes were used in a multiplex reaction. *T. gondii* 529 bp Repeat Element (RE): *forward:* 5′-CACAGAAGGGACAGAAGTCGAA-3′; *reverse:* 5′-CAGTCCTGATATCTCTCCTCCAAGA-3′; *probe:* 5′-CTACAGACGCGATGCC-3′ (IDT). This was normalized to mouse beta-actin using the validated probe: Mm02619580_g (Thermo Fisher Scientific).

### 16S amplicon sequencing

The contents from the small intestine, cecum and large intestine were homogenized in 0.175 mL of water per 50 mg of sample, aliquoted, and stored at -80°C. DNA was extracted from the intestinal contents using the Qiagen DNeasy PowerSoil Pro Kit (cat #47016) per manufacturer’s instructions. 16S ribosomal DNA sequencing was performed by Novogene. Briefly, the V4 hypervariable region of the 16S rRNA gene was amplified using barcoded primers 515F (GTGCCAGCMGCCGCGGTAA) and 806R (GGACTACHVGGGTWTCTAAT) (42). Amplicons were visualized on a 2% agarose gel, quantified with real-time PCR, and pooled in equimolar concentration followed by end-repairing, A-tailing, and Illumina adapters. Libraries were sequenced on an Illumina NovaSeq 6000 platform to generate 250 bp paired-end raw reads. Two negative controls were included during sample processing and sequencing.

### 16S bioinformatics analysis

The 16S sequence with barcode and primer removed were analyzed with DADA2 Workflow for Big Data (https://benjjneb.github.io/dada2/bigdata.html) and dada2 (v 1.26.0) (43). Forward and reverse reads were trimmed using lengths of 150 bp, filtered to contain no ambiguous bases, and contained a minimum quality score of 2. Reads were assembled and chimeras were removed per dada2 protocol.

Taxonomy was assigned to each amplicon sequence variant (ASV) using a combination of the SILVA v128 database (44) and the RDP naïve Bayesian classifier as implemented in the dada2 R package (43). Read counts for ASVs assigned to the same taxonomy were summed for each sample. Alpha diversity of microbiome samples was evaluated using the Shannon alpha diversity index. Beta diversity of the microbiome was measured using Bray-Curtis dissimilarity measures on the basis of relative abundance data. The statistical significance of alpha diversity was determined by pairwise comparisons using Wilcoxon rank sum test with continuity correction, while the significance of beta diversity and relative abundance of each family were measured by permutational multivariate analysis of variance (PERMANOVA), and unpaired *t*-test, respectively.

### Mmvec

For each intestinal content and infectious stage, a table of all detected metabolites and of the bacterial families with abundance >5% were subjected to microbe-metabolite vectors analysis (mmvec) (45), producing conditional probabilities between metabolites and microbiome families. In R (RStudio, version 2023.03.0), non-annotated metabolites and metabolites with mean decrease accuracy lower than 1 from random forest analysis were subsequently excluded from the conditionals file. The subsetted conditionals file was then subjected to Principal Component Analysis (PCA) and visualized using the pca3d package (version 0.10.2) in R.

### MicrobeMASST

To identify a putative microbial origin for metabolites perturbed by infection, MicrobeMASST, an open-source tool integrated into the Global Natural Products Social Molecular Networking (GNPS) platform was used. MicrobeMASST allows the comparison of experimental tandem mass spectrometry (MS/MS) spectra against a comprehensive GNPS database, which contains spectra derived from bacterial, fungal, and archaeal monocultures (46). For this analysis, MS/MS spectra were submitted to the MicrobeMASST interface. The tool performed a spectral similarity search, comparing each input spectrum to the GNPS repository to identify potential microbial sources. The search results provided a list of microbial strains that produced metabolites with similar MS/MS spectral profiles to those detected in our dataset. All spectra were processed using default parameters in MicrobeMASST, except for the following: *m/z* tolerance=0.02, precursor *m/z* tolerance= 0.2 and min matched signals=4.

### Data availability

Metabolomics data were uploaded to MassIVE (**<u>massive.ucsd.edu</u>**), accession number MSV000095605. Molecular networking jobs are accessible through these links: https://gnps.ucsd.edu/ProteoSAFe/status.jsp?task=eaa0f5e4ac6649099029626861b8d843 (feature-based molecular network) and https://gnps.ucsd.edu/ProteoSAFe/status.jsp?task=9391d20cc6a64315994fc21142fd28a1 (MolNetEnhancer). Representative code used to generate manuscript figures has been deposited at https://github.com/CEmiddleton/McCall-lab/tree/main/Toxoplasma. 16S ribosomal RNA sequencing metagenomic data were uploaded to GenBank (NCBI) and are publicly available using accession number PRJNA1198079.

## Results

### Localized metabolic impact of *T. gondii* infection is not explained by local tissue parasite burden

Infectious disease processes are highly localized, with differential sensitivity to pathogen colonization, damage and recovery processes depending on the organ and tissue (47). Prior studies evaluating the consequences of *T. gondii* infection on metabolism have focused on in vitro culture, or in vivo analysis of serum or single tissue sampling sites (16, 17)(9, 11, 16)(48)(49). We therefore sought to generate a systematic understanding of spatial metabolic alterations during acute and chronic *T. gondii* infection. Spatial metabolomic analysis (“chemical cartography”) (50) was performed on eleven sampling sites, selected using the following rationale. Cardiac muscle (site 1) and skeletal muscle (site 2) are major sites of cyst burden during chronic infection. In addition, skeletal muscle (site 2) and liver (site 3) are central metabolic tissues which progressively waste during *T. gondii* infection-induced chronic cachexia (15, 51). During acute *T. gondii* infection, parasites and gut commensal bacteria translocate to the liver, driving Toll-like receptor activation and inflammation (52–54). *T. gondii* interacts with the duodenum (site 4), jejunum (site 5) and ileum (site 6) in the first days post infection, with highest burdens in the distal small intestine (55). Although *T. gondii* has not been documented to infect the cecum (site 7) or large intestine (site 8), these organs support the largest population and diversity of gut commensal bacteria (56). Systemic inflammation can shift gut microbial populations and their metabolic output (57). In addition, we sampled the contents of the small intestine (site 9), cecum (site 10) and large intestine (site 11) to determine if metabolites identified in host tissues were unique or related to any of these gut microbial niches. Each site was sampled in acute (15 days post infection) and chronic (50 days post infection) infection in C57BL/6 mice infected with *T. gondii* and compared to uninfected littermate tissues harvested on the same day (Fig. S1).

In the acute stage of infection, the greatest degree of metabolic alteration was in the large intestine contents (p=0.002, pseudo-F= 9.165), liver (p=0.001, pseudo-F= 6.148), cecum contents (p=0.001, pseudo-F= 4.205) and heart (p=0.001, pseudo-F= 3.563), followed by the cecum (p=0.002, pseudo-F= 2.650), and duodenum (p= 0.011, pseudo-F= 2.455) (Fig. 1A). This pattern was unexpected, since some of the tissues with significant metabolic perturbation, including the colon and cecum, are not expected to interface directly with *T. gondii*. Parasite burden at these sites at 15 days post-infection was low but detectable in most samples (Fig. 1B, Fig. S2). At 15 days post infection, parasite genomic DNA was highest in the skeletal muscle, ileum and jejunum with low but detectable levels in the duodenum, heart and large intestine (Fig. 1B, Fig. S2). Strikingly, there was no correlation between the magnitude of metabolite perturbation and the local parasite burden in each organ during the acute stage (Fig. 2A).

The pattern of overall metabolic perturbation was different in the chronic stage compared to the acute stage. The highest metabolic perturbation in the chronic stage was observed in the large intestine contents (p=0.001, pseudo-F= 4.342), large intestine (p=0.012, pseudo-F= 3.998) and cecum (p= 0.002, pseudo-F= 3.417), followed by cecum contents (p= 0.003, pseudo-F= 3.402) (compare Fig. 1C to Fig. 1A). Other sites had re-normalized compared to the acute stage, excluding skeletal muscle and heart where parasite load was high, as expected (Fig. 1C-D). These data established that the major sites of metabolic perturbation are dynamic from 15 to 50 days post-infection, and that the discordance between local parasite burden and the degree of metabolic perturbation observed in the acute stage is also apparent at chronic infection timepoints.

**Fig. 1.**
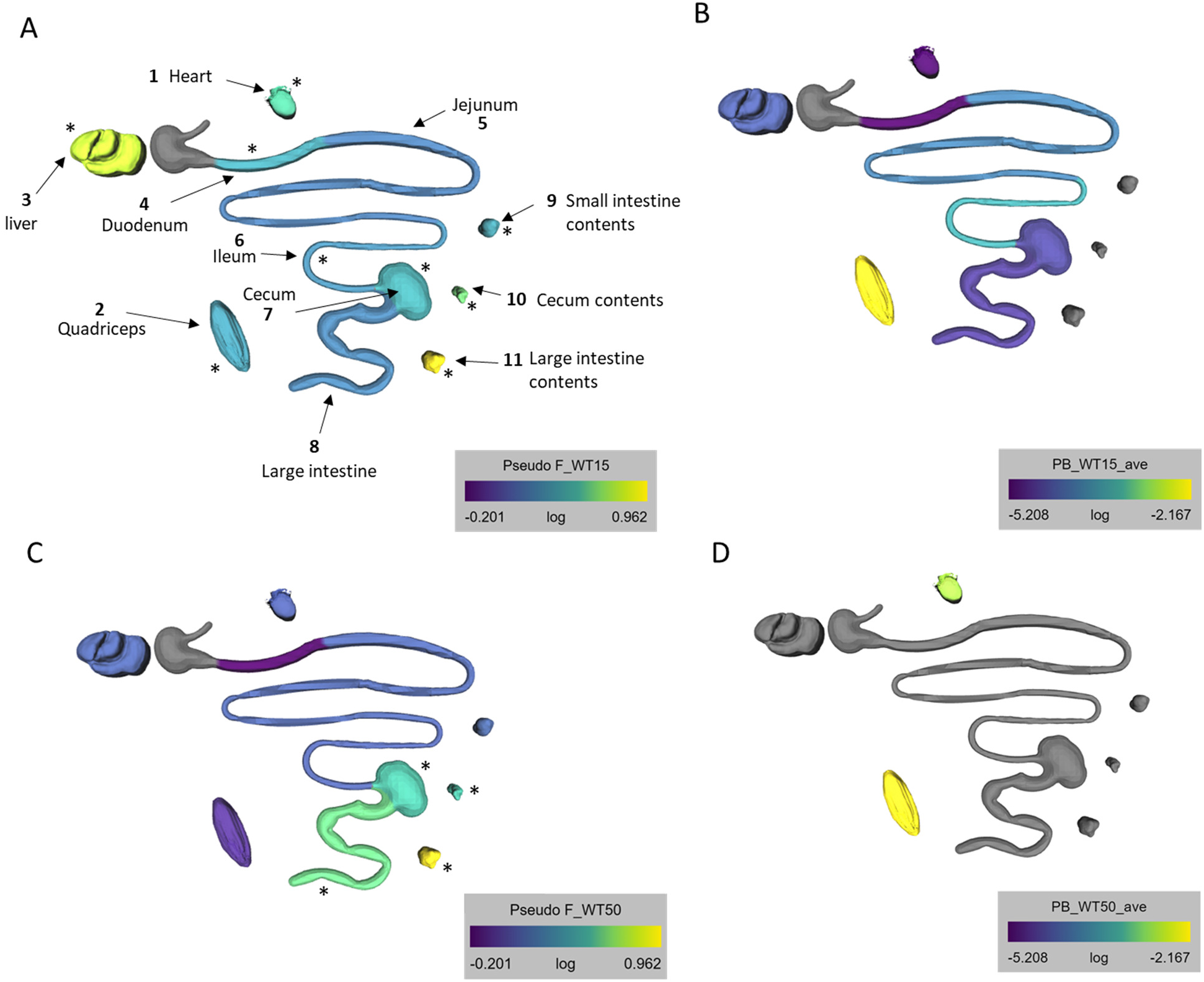
Spatial mismatch between metabolic perturbation and parasite burden at acute and chronic stages of infection. C57BL/6 mice were infected per orally with 20 Me49 *T. gondii* cysts and tissues were dissected at 15 days post infection or (A-B) or 50 days post infection (C-D). The metabolic perturbation in each organ is represented as pseudo-F from pairwise PERMANOVA test between infected and uninfected samples at 15 days post-infection (A) or 50 days post infection (C). The pseudo-F reflects the effect size when comparing infected vs uninfected groups. Its calculation considers within-group distances (infected vs infected or uninfected vs uninfected) and between-group distances (infected vs uninfected), number of samples and number of groups. Average parasite burden in each organ was determined at 15 days post-infection (B) and average parasite load was surveyed in the cardiac muscle and skeletal muscle at chronic infection (D). Gray color denotes organs where metabolites and/or parasite burden were not analyzed. 15 days uninfected N=10, 15 day acute infected N=9, 50 day uninfected N=10, 50 day chronic infected N=7.

**Fig. 2:**
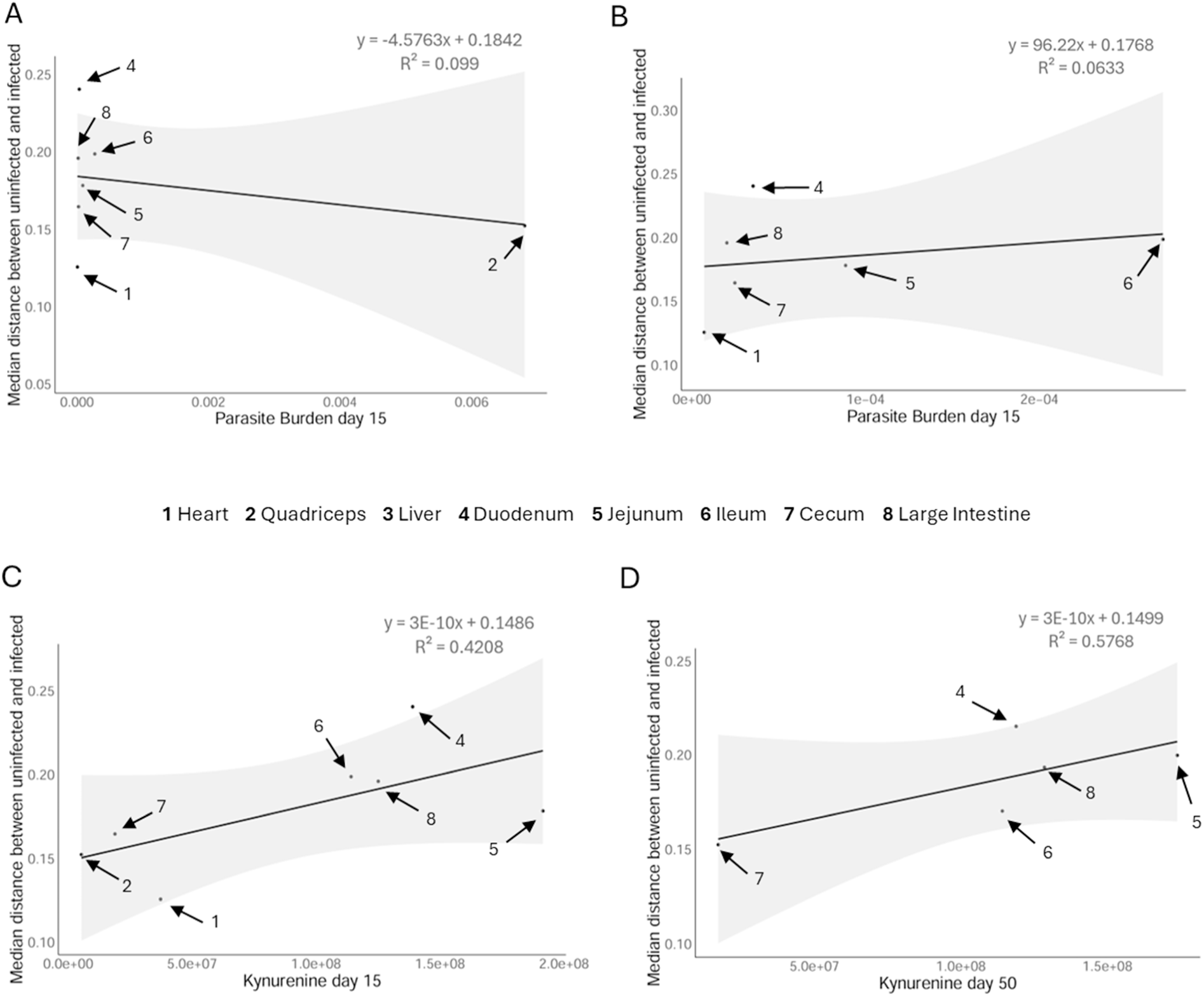
Acute tissue parasite burden and the severity of tissue metabolic perturbation do not correlate, unlike kynurenine levels. (A) Metabolic perturbation represented as median metabolic distance from pairwise PERMANOVA between infected and uninfected samples (Y axis) was plotted relative to the average parasite burden in each organ for each tissue in acute infection. (B) Median metabolic distance relative to average parasite burden at the acute stage with the quadriceps outlier removed. (C) and (D) Median metabolic distance between uninfected and infected tissues relative to median kynurenine levels in acute infection (C) and chronic infection (D). The grey shaded region indicates the confidence interval for the linear model. Organs where kynurenine was only detected in one sample or fewer are excluded from analysis.

To identify metabolites that were differentially regulated in response to acute infection and determine if any were shared between tissues, random forest analysis was applied (Dataset S1). None of the metabolites that changed in response to infection were common across all eleven tissue sampling sites (Fig. 3), even though, as expected, most metabolite features (irrespective of impact of infection) were shared across multiple or even most sampling sites (Fig. S3). Dominant patterns of overlap in infection-perturbed metabolites were between sampling sites expected to share common biology, such as cecum contents and large intestine contents, small intestine contents and cecum contents, jejunum and ileum, and large intestine and cecum (Fig. 3, Dataset S1). However, commonalities were also observed between liver and heart (15 exclusively overlapping differential features at 15 days post-infection; 10 at 50 days post-infection), liver and quadriceps (12 exclusively overlapping differential features at 15 days and 50 days post-infection) and heart and quadriceps (12 exclusively overlapping differential features at 15 days and 50 days post-infection), possibly reflecting common patterns across sites of high parasite burden, muscular tissue in general, or sites implicated in *Toxoplasma* cachexia (15, 51) (Fig. 3).

Lipids were correlated with parasite burden. Ceramides negatively correlated with parasite burden in the large intestine. Ceramides regulate cell signaling pathways (58). Ceramide biosynthesis in *T. gondii* has been linked to the parasite’s ability to adapt to different host environments, therefore this metabolite could be parasite derived (59). Glycerophosphocholines positively correlated with parasite burden in the heart and ileum in the acute stage, and glycerophosphocholines negatively correlated with parasite burden in the ileum and jejunum in the acute stage. Abnormal phosphatidylcholine ratios could influence energy metabolism and can be a sign of disease progression in metabolic disorders including atherosclerosis, and obesity (60). However, none of these metabolite-parasite burden correlations were significant after correction for multiple hypothesis testing.

**Fig. 3.**
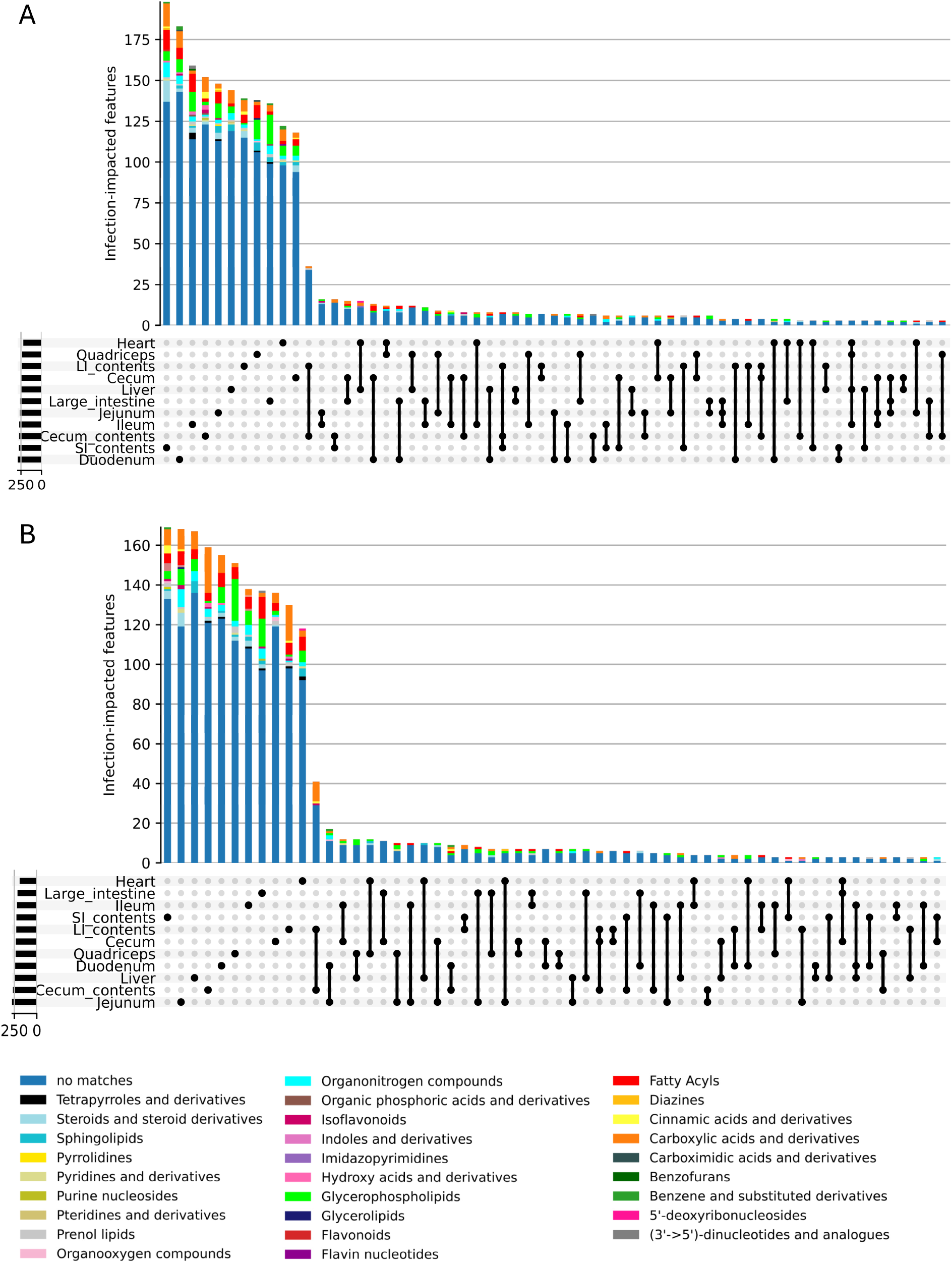
Tissue-specific impact of infection. Carboxylic acids, fatty acyls, glycerophospholipids, and organonitrogen compounds are perturbed during *T. gondii* infection in all organs, although individual metabolites are rarely differentially enriched in multiple tissues. UpSet plots representing the intersection of perturbed metabolites at 15 days post infection (A) or 50 days post infection (B). The bars indicate the number of perturbed metabolites in a specific organ only (left part of the graph, tissue indicated by single black dot below the bars) or commonly perturbed between two or more organs (right part of the graphs, pairwise comparisons indicated by black dots and connecting line below the bars). Only intersections between 3 or more sampling sites are displayed. The chemical families of each perturbed metabolite are shown by the colors in the bar graphs.

### Magnitude of localized metabolic impact of *T. gondii* infection is partially explained by local tissue inflammation

Having established that shifts in metabolic homeostasis are not strongly explained by concurrent, local *T. gondii* parasite burden, we next assessed whether tissue-specific pro-inflammatory immune responses could be responsible for the observed localized metabolic perturbations. Immune responses can reshape local metabolism, and local immune responses can remain activated even after pathogen clearance (61, 62). Kynurenine is induced by pro-inflammatory signaling and regulates immunity (63). Because we could readily detect it by mass spectrometry in our dataset, we used tissue kynurenine levels as a surrogate for the degree of local pro-inflammatory signaling (*m/z* 209.091 RT 0.47 min ([M+H]^+^ adduct) and *m/z* 192.065 RT 0.48 min ([M+H-NH3]^+^ adduct)). Kynurenine annotation was previously validated using pure standards under the same instrumental conditions, at a similar retention time (level 1 annotation confidence according to the metabolomics standards initiative (35)(23)). Kynurenine was significantly increased by infection in 7 sampling sites during the acute stage. We observed a moderate correlation between the metabolic impact of infection and kynurenine levels in the acute stage (R^2^= 0.4208), with a stronger correlation in the chronic stage of infection (R^2^=0.5768) (Fig. 2C-D). In contrast to this strong relationship between kynurenine and between-organ differences, kynurenine did not correlate with within-organ differences between mice (Fig. S4).

Correlating infection-impacted metabolite features (as determined by random forest analysis) to kynurenine levels in each organ showed differential patterns between sampling sites (Dataset S1). Overall, 2.97% to 50% of infection-impacted features were correlated with kynurenine, with the lowest of 2.97% in the quadriceps in the chronic stage and the highest of 50% in the ileum in the acute stage. There were a few common metabolites correlated with kynurenine and with parasite burden, including ile-leu in the duodenum, propionylcarnitine in the heart, and adenosine and multiple phospholipids in the ileum, all in the acute stage (uncorrected Spearman correlation p-value p<0.05).

Metabolites related to inflammation pathways were affected in both acute and chronic stages of infection and correlated with kynurenine levels. Lysophosphatidylcholine (LPC) originates from the cleavage of phosphatidylcholine by phospholipase A2. LPC can contribute to inflammation by activating ion channels and increasing the release of inflammatory factors (64). Metabolites annotated as Lyso-PC(16:0) and Lyso-PC (22:6) were highly correlated (Spearman correlation coefficient > 0.5, uncorrected p<0.05) with kynurenine in the ileum in the acute stage of infection. The hemin cation was highly correlated with kynurenine in the large intestine, ileum and jejunum in the acute stage of infection. Hemin is an iron containing metalloporphyrin which activates the heme oxygenase (HO) enzyme with cardioprotective and anti-inflammatory effects (65). In the acute stage, the heart was found to have increased xanthine levels, positively correlated with kynurenine levels (Spearman uncorrected p-value p=2.79E-05). Xanthine is a crucial intermediate in the degradation of purine nucleotides, leading to the production of uric acid. This pathway has reactive oxygen species (ROS) as byproducts, which play an important role in the progression of inflammatory disorders (66)(67). Reduced glutathione and S-hexyl-glutathione were negatively correlated with kynurenine in the heart in acute infection (uncorrected Spearman correlation p<0.05). These metabolites are mostly involved in oxidative stress defense. Their negative correlation with kynurenine could suggest antioxidant usage in response to inflammation. Beta oxidation related metabolites, specifically acylcarnitines, were also strongly correlated with kynurenine. For example, acetyl-carnitine was elevated in the heart, while propionylcarnitine and carnitine were negatively correlated with kynurenine in the heart. Propionylcarnitine was likewise negatively correlated with kynurenine in the jejunum during acute infection (uncorrected Spearman correlation p-value p<0.05).

### Role of the gut microbiome in local tissue metabolic responses to infection

Immune responses and the microbiome can influence each other, but the microbiome can also have effects independent of immune alterations, particularly through small molecule signals (68). In this study, we observed significant changes in overall metabolism in cecum and colon contents during acute and chronic infection, even though parasite burden was low at these sites (Fig. 1). The cecum and the colon are major sites of microbiome metabolic activity (69)(68).

To understand how shifts in commensal microbiota are linked to metabolite levels, we performed 16S sequencing of small intestine, cecum and large intestine contents (Fig. 4, Fig. S5, Fig. S6). Bray non-metric multidimensional scaling (NNMDS) analysis showed that the microbial 16S composition in the small intestine, cecum and colon contents were distinct in uninfected and infected littermates in acute infection (Fig. 4A). The disruption in the microbial composition between infected and uninfected mice remained significant in chronic infection (Fig. 4B). At acute infection, increased *Peptostreptococcaceae* (Fig. 4C) and *Clostridiaceae* (Fig. 4D) and a decrease in *Bifidobacteriaceae* (Fig. 4E) were observed in the small intestine and large intestine contents in acute infection. In addition, there were some infection-dependent changes that were unique to each intestinal tract site. For example, there was a significant increase in small intestine *Erysipelotrichaceae* and a decrease in *Muribaculaceae*, (Fig. 4F-G). The cecum had an increase in *Bacteroidaceae* (Fig. 4H). In chronic infection, there were still microbial species that were significantly differentially enriched in the small intestine contents, including *Bifidobacteriaceae (*also in the colon contents) (Fig. 4I), *Erysipelotrichaceae* (Fig. 4J) and *Desulfovibrionaceae* (Fig. 4K). In addition, the increase in large intestinal contents *Muribaculaceae* was still observed at chronic infection (Fig. 4L).

**Fig. 4.**
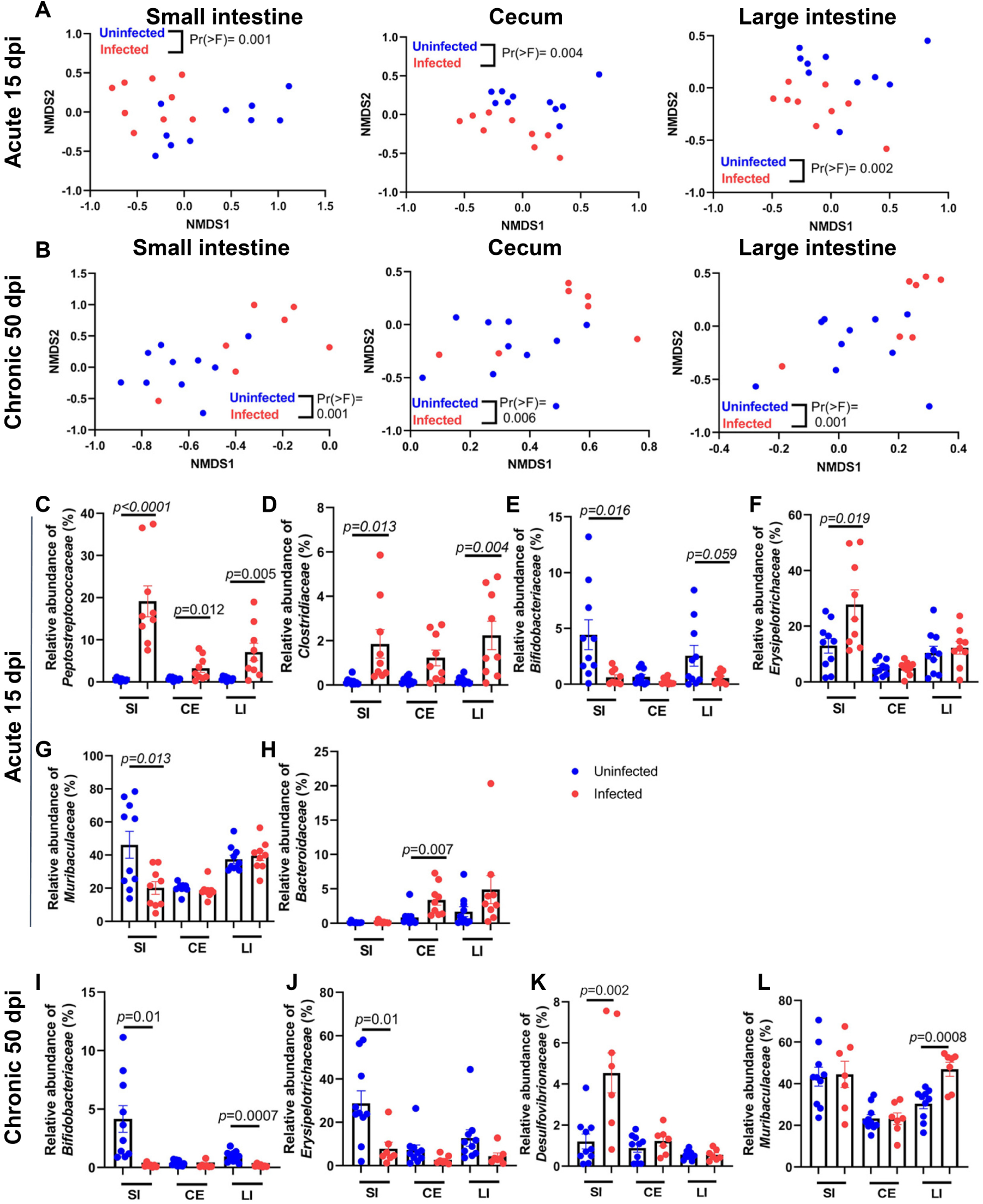
Impact of acute and chronic *T. gondii* infection on the microbiome. (A-B) Beta diversity of bacterial components in the small intestine, cecum, and large intestine contents was compared using the Brays NMDS analysis of 16S rRNA sequencing at 15 days post infection (A) or 50 days post infection (B). Statistical significance was determined by Permutational multivariate analysis of variance (PERMANOVA). (C-H) At 15 days post infection, the relative abundance of (C) *Peptostreptococcaceae*, (D) *Clostridiaceae,* (E) *Bifidobacteriaceae*, (F) *Erysipelotrichaceae*, (G) *Muribaculaceae*, and (H) *Bacteroidaceae* in the small intestine (SI), cecum (CE) and large intestine (LI) contents from uninfected and infected mice. (I-L) At 50 days post infection, the relative abundance of (I) *Bifidobacteriaceae,* (J) *Erysipelotrichaceae*, (K) *Desulfovibrionaceae*, and (L) *Muribaculaceae* in the fecal contents from uninfected and infected mice. Error bars represent the mean +/-SEM. N=9-10 per group.

These microbiome compositional changes may be shaping the metabolic response to infection. Alternatively, changes in luminal nutrient availability during infection could be causing the observed effects of infection on the microbiome. To further explore these relationships, we performed microbe-metabolite vectors (mmvec) analysis (45) on the most abundant microbial families and the metabolite peak areas, filtered to those identified as perturbed by infection using random forest analysis. Multiple patterns of co-occurrence were observed, such as between steroids and *Clostridiaceae* family members, *Erysipelotrichaceae* family members and *Peptostreptococcaceae* family members in cecum contents during acute infection or with *Muribaculaceae* and *Staphylococcaceae* (lower in infected versus uninfected p=0.02) family members in the large intestine contents during acute infection. In the small intestine contents, *Lachnospiraceae* and *Muribaculaceae* showed opposite occurrence patterns to organonitrogen compounds during chronic infection (Fig. 5). Of these co-occurrence patterns, only *Peptostreptococcaceae* were differentially abundant with infection, being significantly increased by acute infection in the cecum contents. These results highlight how these metabolic patterns may be predominantly driven by changes in microbial metabolism and microbial gene expression rather than differential microbial abundance.

As further support of a possible microbial origin for these metabolites, we performed microbeMASST analysis. MicrobeMASST is an open source tool within the GNPS system that matches experimental tandem MS spectra to spectra within the GNPS repository which were retrieved from bacterial, fungal and archaeal monocultures (46). Overall, many infection-impacted features had a candidate microbial origin. For example, the cecum contents in the chronic stage had the highest percentage of microbeMASST matches (62.7% matches), and the main chemical families with matches were phospholipids, amino acids and peptides. The lowest percentage of microbeMASST matches was observed in the liver in the acute stage of infection (45.5% matches).

Corresponding findings were also observed for several features in the cecum, large intestine and small intestine contents between mmvec and microbeMASST analysis. For example, in the cecum contents at day 15, *m/z* 785.589 retention time (RT) 2.78 min had a co-occurrence with *Peptostreptococcaceae* which was supported with a microbeMASST match to *Peptostreptococcus* genus. In the large intestine contents at day 50, for metabolite feature *m/z* 187.144 RT 0.45 min, mmvec found an *Akkermansiaceae* co-occurrence and microbeMASST found a match to *Akkermansia muciniphila*. In the small intestine contents at day 15, metabolite feature *m/z* 175.118 RT 0.26 min had an mmvec match to *Lactobacillaceae* family, and microbeMASST had several matches to genera within this family. At day 50, for metabolite feature *m/z* 205.096 RT 3.47 minutes, mmvec found co-occurrence with *Micrococcaceae,* supported by a microbeMASST matches to *Micrococcus luteus*. The consistent results from both microbeMASST and mmvec analyses support a microbial origin for these infection-impacted metabolites.

**Fig. 5.**
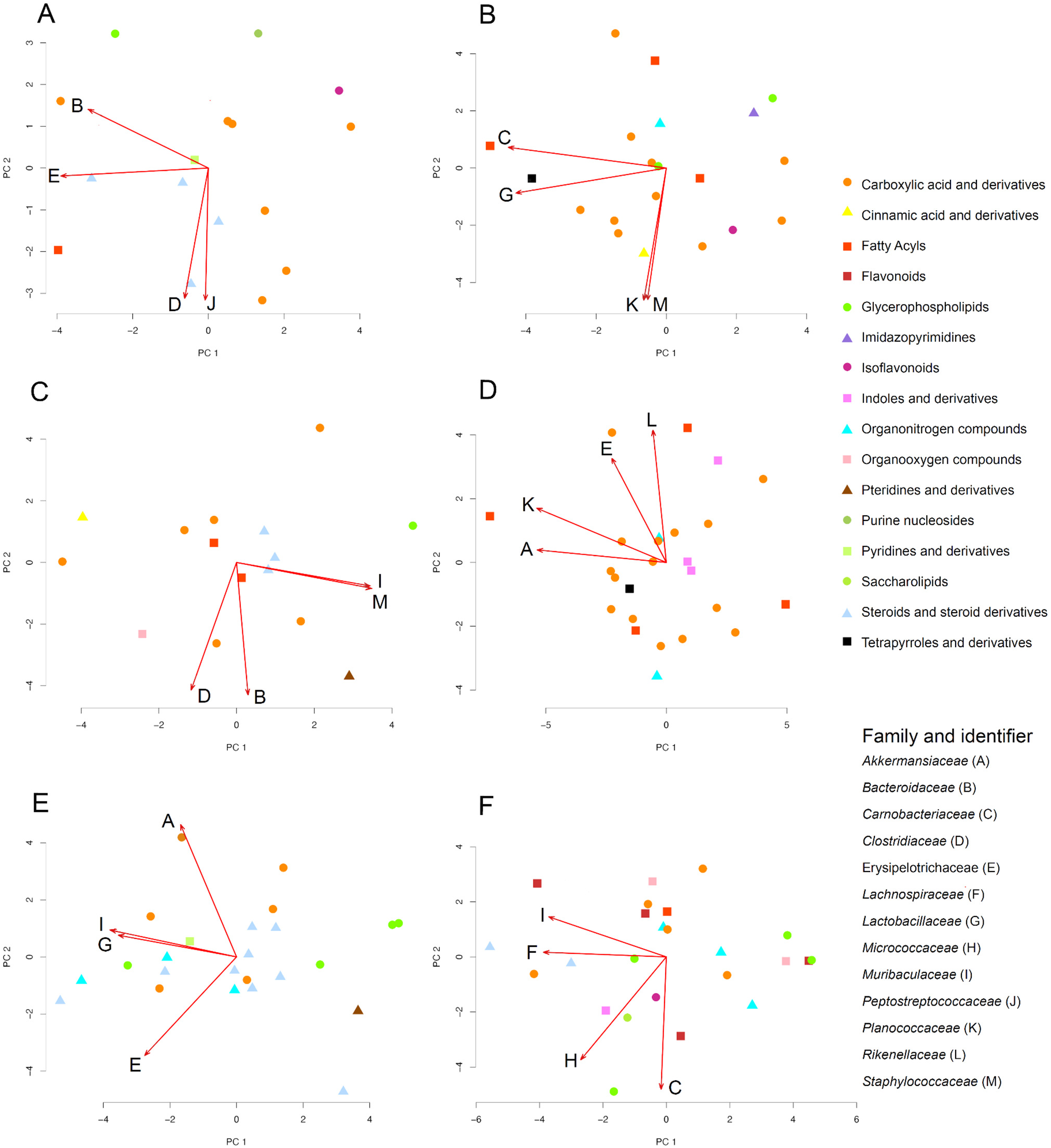
Co-occurrence between the metabolites most perturbed by infection and gut microbiome families in GI tract contents. mmvec analysis was applied to the most abundant microbial families (>5% abundance) and on the metabolite peak areas, followed by PCA analysis. Only metabolites with a random forest mean decrease accuracy >1 and with annotations are displayed. (A) Cecum contents in the acute stage. (B) Cecum contents in the chronic stage. (C) Large intestine contents in the acute stage. (D) Large intestine contents in the chronic stage. (E) Small intestine contents in the acute stage. (F) Small intestine contents in the chronic stage. Red arrows, 4 most influential microbiome families. Capital letters, identifiers for each microbiome family.

## Discussion

Metabolic changes in association with infectious diseases have been extensively studied (70)(71)(72), and demonstrated to play a causal role in disease progression (e.g. (73)(25)(74)). Pathogen-derived metabolites, damage-induced signals, microbiome-derived molecules and immune-mediated metabolic responses have all been implicated as regulators of these adaptive and maladaptive metabolic changes (8). However, their relative importance across organs in driving overall metabolic changes is not well characterized. Here, using *T. gondii* infection as a model, we assessed the relative contribution of tissue site, local tissue parasite burden, local tissue immune responses, infection duration, and microbiome composition, in regulating tissue metabolic responses. We observed a limited contribution of local tissue parasite burden (Fig. 1, Fig. 2), though this does not preclude prior parasite colonization, cleared at the time of sample collection, from having caused the observed metabolic changes. This observation concurs with our findings with the parasites *Trypanosoma cruzi* (*25, 61, 75*) and *Leishmania major* (*76*), and in SARS-CoV-2 infection (77). In contrast, in a mouse model of acute influenza A virus infection, a positive correlation was observed between metabolic perturbation and tissue viral load (23).

Unlike tissue parasite burden, tissue inflammation was positively correlated with metabolic impacts of infection (Fig. 2BC), suggesting a dominant regulatory role of inflammation over direct parasite effects. Immunity and metabolism are tightly interlinked (78). Indeed, many of the infection-impacted metabolites correlated with kynurenine are known to intersect with immunity, including lysophosphatidylcholines in the ileum during acute infection, hemin in the large intestine, ileum and jejunum in the acute stage of infection, and xanthine, reduced glutathione and S-Hexyl-Glutathione in the heart in acute infection. LPC can increase chemokines which attract inflammatory cells and increase the release of inflammatory factors such as IFN-y (64). Hemin is an activator of the antiviral, antioxidant, and anti-inflammatory heme oxygenase-1 (HO-1) enzyme (79). HO-1 provides cytoprotection against inflammatory and oxidative injury (80). In COVID-19 viral infection, the HO-1 enzyme exerts antiviral properties by interfering with the interferon (IFN) pathway (80). The glutathione system is a robust antioxidant system in the heart responsible for scavenging ROS (66, 81). Decreased levels indicate antioxidant usage corresponding to increased inflammation and could indicate cardiovascular damage (82). ROS production is significant in *T. gondii* infection (82, 83).

The large intestine contents, large intestine and cecum were the most metabolically perturbed tissue sites during chronic infection (Fig. 1C). Per oral infection with *T. gondii* leads to reduction in commensal diversity during acute infection and outgrowth of Gram-negative species associated with lethal ileitis (84)(52, 84). Non-lethal infection has shown that dysbiosis persists into chronic infection, after inflammatory cytokines and tissue pathology in the small intestine has returned to baseline levels (19, 85). Similar to previous reports, we found that several microbial families associated with pathology were enriched in acute and/or chronic infection including (19, 52, 86). However, previous studies have exclusively evaluated fecal microbes. Our assessment of the small intestine and cecum contents revealed a new role for the anaerobic, Gram-positive family *Peptostreptococcaceae* which were more highly enriched in the small intestine contents than other sites (Fig. 4C). *Peptostreptococcaceae* and *Clostrridiaceae* (Fig. 4C-D) have been associated with frailty and muscle wasting in aged human participants in the BIOSPHERE study (87). Unlike the BIOSPHERE study, we found a reduction in the abundance in *Bifidobacteriaceae* in the small and large intestine contents at acute infection (Fig. 4E). This anaerobic Gram-positive population is involved in oligosaccharide metabolism and has been explored as a probiotic treatment for ulcerative colitis and inflammatory bowel syndrome (88, 89). As expected, the microbial populations were divergent in controls associated from 15 days post infection and 50 days post infection populations, which may be related to cohort and/or age-related differences in each cohort (Fig. S5). Despite this natural variation, *Erysipelotrichaceae* was found to be increased in the small intestine during acute infection (Fig. 4F) but significantly depressed during chronic infection (Fig. 4J) compared to uninfected controls. This family includes highly immunogenic species which are positively correlated with dietary fat intake and obesity (90–92). A possible microbiome source for multiple metabolites impacted by infection was identified in this study. The carboxylic acids chemical family in particular had strong co-occurrence with influential microbe families in all three gastrointestinal organ contents at both acute and chronic stages. The availability of amino acids affects gut microbiome growth, and the gut microbiome affects amino acids absorption and metabolism (93–95). Amino acid auxotrophies of *T. gondii* can be another reason for amino acid perturbation during toxoplasmosis (96).

In summary, our study provides valuable insights into the metabolic perturbations induced by *T. gondii* infection across organs and potential mechanisms underlying the observed changes. Future research should focus on elucidating the functional consequences of these perturbations, further exploring their relationship with immune responses, and investigating the effect of metabolic alteration on the gut microbiome and downstream changes in host metabolism during *T. gondii* infection.

## Acknowledgements

LIM holds an Investigators in the Pathogenesis of Infectious Disease Award from the Burroughs Wellcome Fund. This work was supported by NIH/NIGMS R35GM138381 (SEE).

