## Supplemental figures and tables for "Spatially-divergent metabolic impact of experimental toxoplasmosis: immunological and microbial correlates"

**Supporting information:**

**Table S1. MZmine parameters (MZmine2 version 2.53).**

| | | |
|---|---|---|
| Mass Detection | MS1 Noise Level | 4.0E5 |
| | MS2 Noise Level | 1.00E+03 |
| | Mass Detector | Centroid |
| ADAP Chromatogram Builder | Min group size in # of scans | 5 |
| | Group intensity threshold | 4.0E5 |
| | Min highest intensity | 1.2E6 |
| | *m/z* tolerance | 0.001 *m/z* or 10 ppm |
| Chromatogram Deconvolution: local minima algorithm | Chromatographic threshold | 20 |
| | Search minimum in RT range (min) | 0.2 |
| | Minimum relative height | 26 |
| | Minimum absolute height | 1.6E6 |
| | Min ratio of peak top/edge | 1.0 |
| | Peak duration range (min) | 0.01-1.00 |
| | *m/z* Range for MS2 Scan Pairing (Da) | 0.01 |
| | RT Range for MS2 Scan Pairing (min) | 0.1 |

| | | |
|---|---|---|
| Isotopic Peak Grouper | Retention Time Tolerance (min) | 0.1 |
| | *m/z* tolerance (ppm) | 10 |
| | Monotonic Shape | Yes |
| | Maximum Charge | 3 |
| | Representative isotope | Lowest *m/z* |
| Join aligner | *m/z* tolerance (ppm) | 10 |
| | *m/z* to RT weight | 10000 to 10 |
| | Retention Time Tolerance (min) | 0.32 |
| Row filtering | Retention Time | 0.20-7 min |
| | Remove previous peak list | Disabled |
| | Keep only peaks with MS2 scan | Enabled |
| | Minimum peaks in a row | 5 |

**Dataset S1. Infection-impacted metabolites and their correlation to parasite burden and kynurenine.**

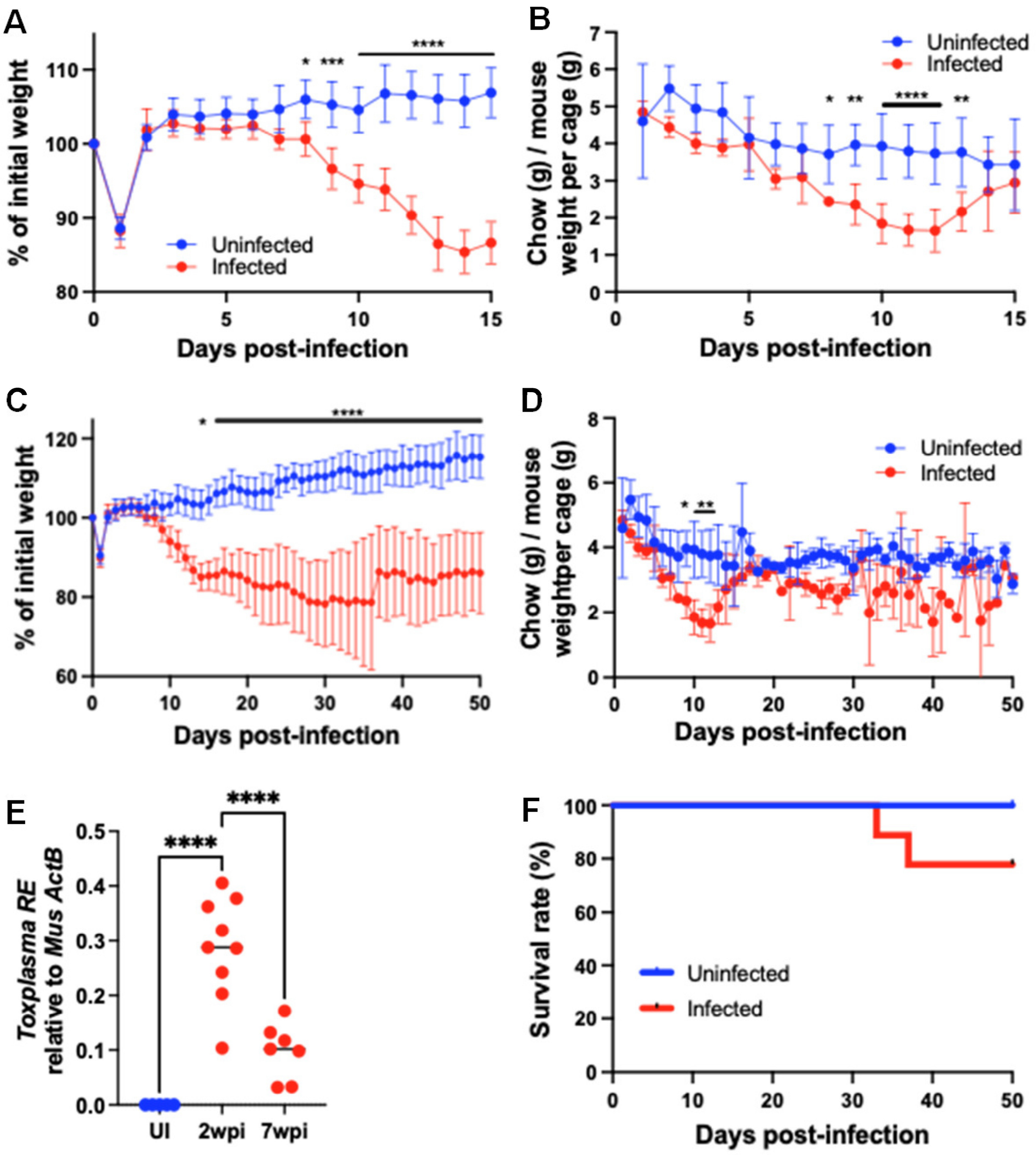


**Fig. S1. Impact of infection on mouse weight and food intake.** C57BL/6J mice were orally infected with 25 Me49gLuc *T. gondii* cysts of the Me49 strain or mock injected with PBS. (A, C) Change in body weight normalized one day before fasting and per oral infection. (B, D) 24 hour food intake per cage normalized to the pooled weight of all mice in the cage. Unpaired Student's t-test with Sidak method to correct for multiple comparisons. Error bars represent standard error mean. (E) *T. gondii* RE levels relative to host beta-actin in brain genomic DNA at 15 days or 50 days post-infection. N=10 uninfected mice and N=7-9 infected mice per group pooled from two independent experiments. Statistical significance was determined by One-way ANOVA. (F) Survival curves of mice represented in (C).

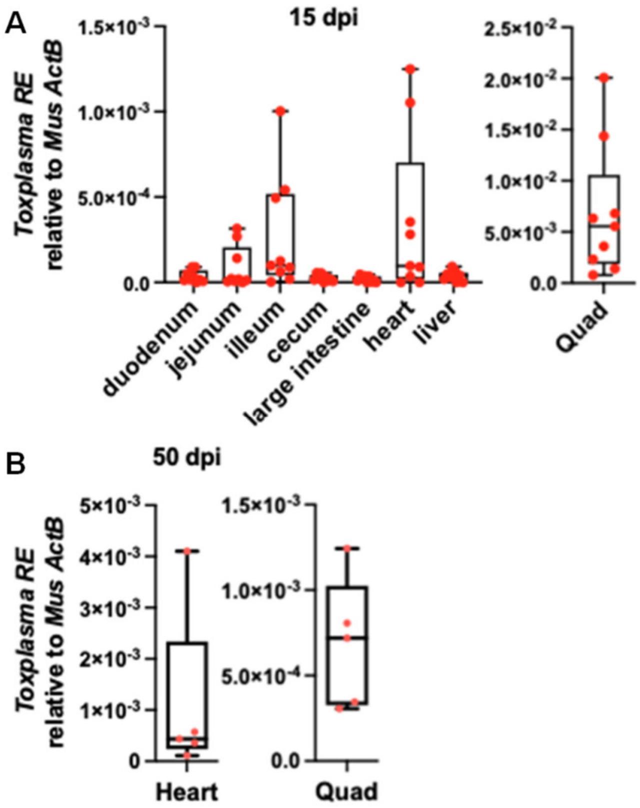


**Fig. S2. Parasite burden in the small intestine duodenum, jejunum, ileum, cecum, large intestine, heart, liver and quadriceps (Quad) at 15 (A, N=9) or 50 (B, N=7) days post infection as described in Fig. 1.** Box and whisker plots represent median, first and third quartile.

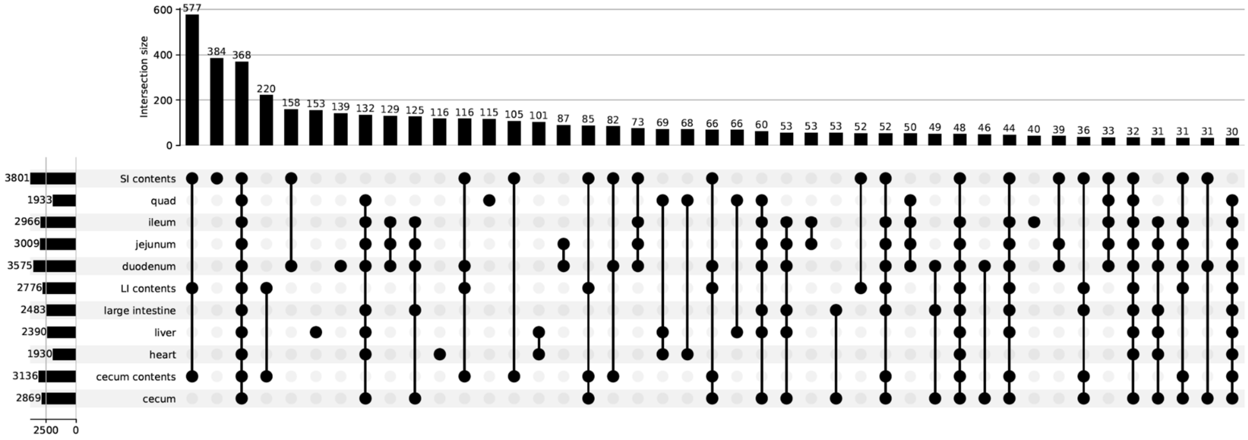


**Fig. S3. Most metabolites are shared across all sampling sites.** All detected metabolites from all groups and all timepoints were analyzed with regards to the sampling sites where they were observed. Unlike infection-impacted metabolites (Fig. 3), considerable overlap of detected metabolites (irrespective of the impact of infection) was observed (connected black dots). Only intersections of 30 or more members are displayed.

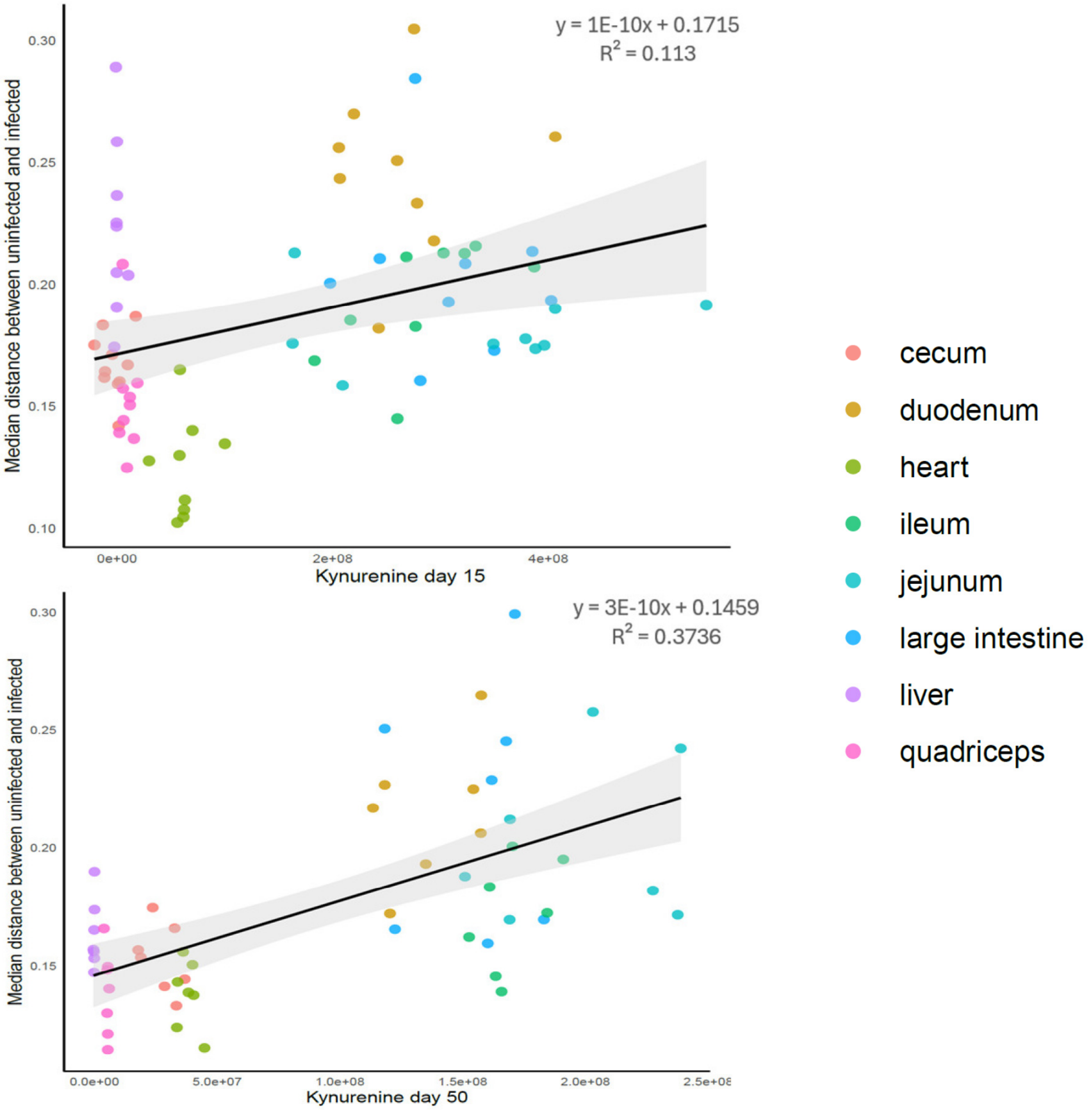


**Fig. S4. Lack of correlation between kynurenine levels and metabolic impact on a per-mouse basis.** Data represent the correlation between Kynurenine levels measured in each infected mouse, colored by tissue type. 15 days post infection N=7, 50 days post infection N=9.

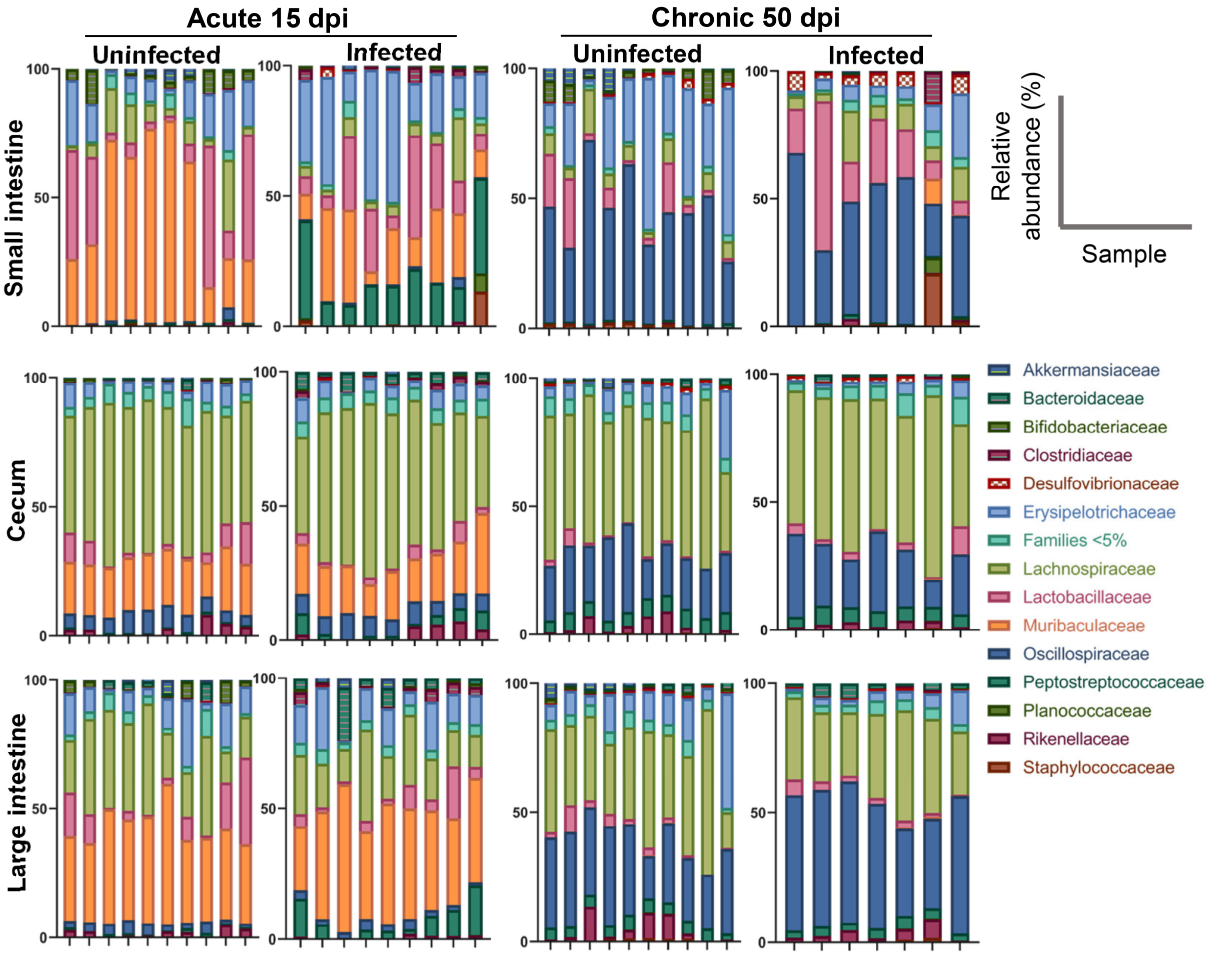


**Fig. S5. The relative abundance of small intestine, cecum and large intestine microbial bacteria from uninfected or *T. gondii*-infected mice.** C57BL/6J mice were orally infected with *T. gondii* or left uninfected. Bacterial composition at the family level in the small intestine, cecum, and large intestine contents were quantified using 16S sequencing in acute (A) and chronic (B) infection.

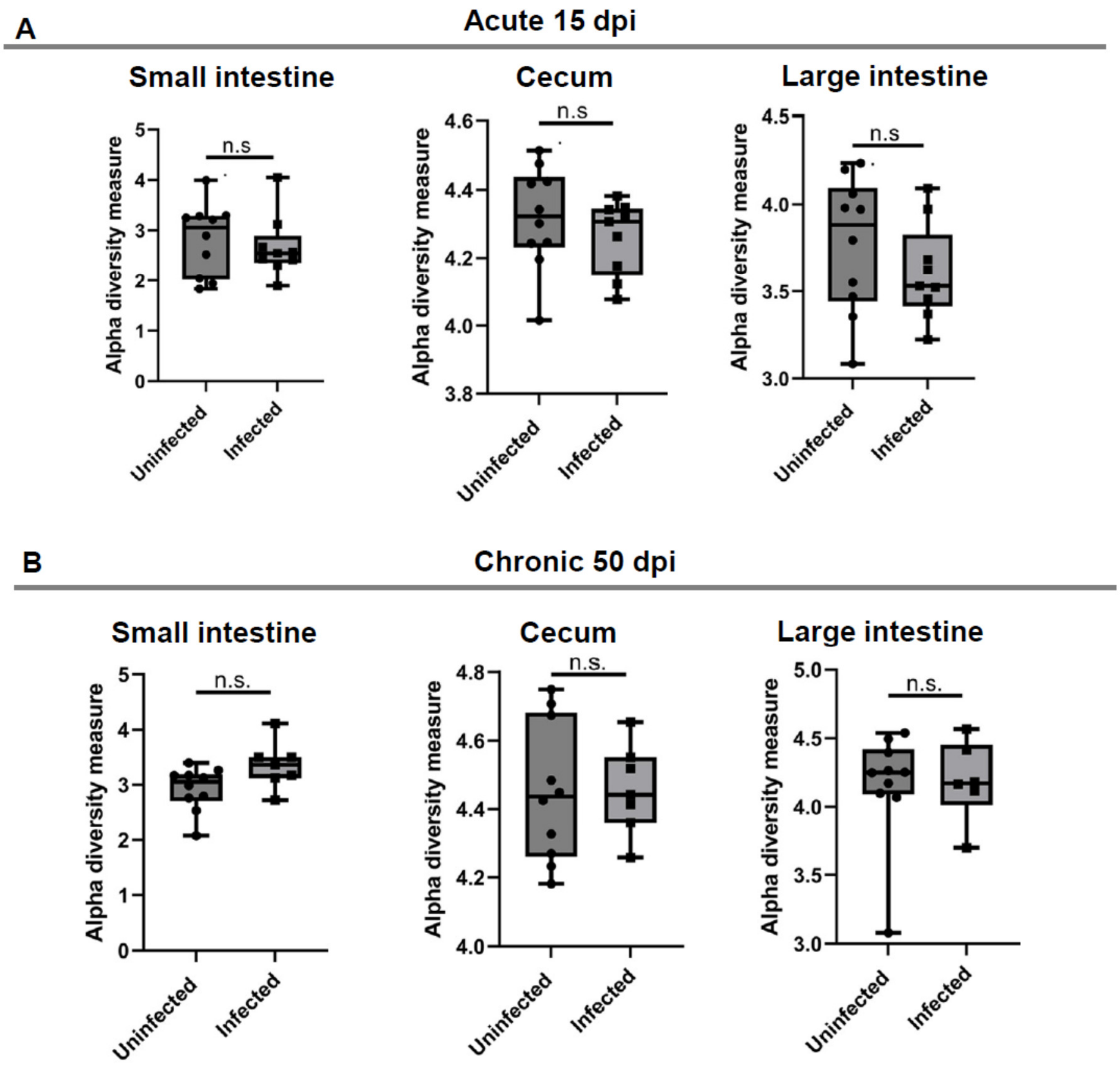


**Fig. S6. *T. gondii* infection does not significantly impact intestinal bacterial alpha-diversity at 15 and 50 days post infection.** 15 or 50 days post-infection, the contents of small intestine, cecum, and large intestine were collected from *T. gondii*-infected or uninfected C57BL/6J mice. The bacterial components in the intestinal contents were analyzed by 16S sequencing. Bacterial alpha diversity in the small intestine, cecum, and large intestine contents was determined using the Shannon index at 15 days post infection (A) or 50 days post infection (B). Max, min and median are presented in each boxplot. Statistical significance was determined by unpaired Student's t-test. Each dot represents an individual mouse. N=10 uninfected, N=9 15 dpi infected, N=7 50 dpi infected.
